# Glucocorticoid Receptor-Regulated Enhancers Play a Central Role in the Gene Regulatory Networks Underlying Drug Addiction

**DOI:** 10.1101/2022.01.12.475507

**Authors:** Sascha H. Duttke, Patricia Montilla-Perez, Max W. Chang, Hairi Li, Hao Chen, Lieselot L. G. Carrette, Giordano de Guglielmo, Olivier George, Abraham A. Palmer, Christopher Benner, Francesca Telese

## Abstract

Substance abuse and addiction represent a significant public health problem that impacts multiple dimensions of society, including healthcare, the economy, and the workforce. In 2021, over 100,000 drug overdose deaths were reported in the US, with an alarming increase in fatalities related to opioids and psychostimulants. Understanding the fundamental gene regulatory mechanisms underlying addiction and related behaviors could facilitate more effective treatments. To explore how repeated drug exposure alters gene regulatory networks in the brain, we combined capped small (cs)RNA-seq, which accurately captures nascent-like initiating transcripts from total RNA, with Hi-C and single nuclei (sn)ATAC-seq. We profiled initiating transcripts in two addiction-related brain regions, the prefrontal cortex (PFC) and the nucleus accumbens (NAc), from rats that were never exposed to drugs or were subjected to prolonged abstinence after oxycodone or cocaine intravenous self-administration (IVSA). Interrogating over 100,000 active transcription start regions (TSRs) revealed that most TSRs had hallmarks of bonafide enhancers and highlighted the KLF/SP1, RFX, and AP1 transcription factors families as central to establishing brain-specific gene regulatory programs. Analysis of rats with addiction-like behaviors versus controls identified addiction-associated repression of transcription at regulatory enhancers recognized by nuclear receptor subfamily 3 group C (NR3C) factors, which include glucocorticoid receptors. Cell-type deconvolution analysis using snATAC-seq uncovered a potential role of glial cells in driving the gene regulatory programs associated with addiction-related phenotypes. These findings highlight the power of advanced transcriptomics methods to provide insight into how addiction perturbs gene regulatory programs in the brain.

## Introduction

Drug addiction and related health problems impact millions of lives in the United States and impose an enormous medical, social, and economic burden on society (Fan et al., 2019). Addiction is a chronic relapsing disorder characterized by diminished control over drug-seeking, compulsive consumption despite negative consequences resulting from drug use, and relapse to drug-taking even after years of abstinence. These enduring effects suggest that chronic drug exposure causes persistent changes in the brain that underlie the development of addiction-related behaviors. The transition from recreational to compulsive drug-seeking is associated with the recruitment of brain reward and stress systems (Koob et al., 2014), including the corticostriatal circuitry that involves the prefrontal cortex (PFC) and the nucleus accumbens (NAc) (Koob and Volkow, 2016). This transition is a critical step in the emergence of compulsivity, which leads to loss of inhibitory control over drug use by recruitment of neuronal populations in the prefrontal cortex (PFC) (Koob and Volkow, 2016).

Numerous studies have demonstrated that long-lasting changes in gene expression patterns in brain regions of the reward pathway are a critical mechanism by which substances of abuse lead to persistent drug-induced neuroadaptations (Russo et al., 2010; Gipson et al., 2014). These neuroadaptations manifest as changes in excitability, synaptic function, and structure, which ultimately contribute to the increased risk of relapse after prolonged abstinence (Dong et al., 2017). It is well known that different drugs of abuse act through distinct receptors but engage convergent pathways that activate or repress the activity of transcriptional factors (TFs) or epigenetic regulators, which in turn drive changes in gene expression patterns (Pierce et al., 2018; Hamilton and Nestler, 2019; Teague and Nestler, 2021; Werner et al., 2021). Numerous studies have elucidated the role of crucial TFs in regulating gene expression patterns altered by repeated exposure to addictive drugs, including opioids and cocaine. These TFs include AMP response element-binding protein (CREB), ΔFOSB, nuclear factor κB (NFκB), early growth response protein 3 (EGR3), and nuclear receptor subfamily 4 group a member 1 (NR4A1) (Hope et al., 1994; Carlezon et al., 1998; Barrot et al., 2002; McClung and Nestler, 2003; Zachariou et al., 2006; Chandra et al., 2015; Carpenter et al., 2020). In parallel, numerous studies have begun to uncover chromatin-mediated mechanisms that contribute to behavioral responses to addictive drugs, such as drug-induced post-translational modification of histone proteins (Stewart et al., 2021).

Despite this knowledge, remarkably little is known about the gene regulatory mechanisms responsible for driving these changes. Mammalian gene expression programs are orchestrated by the collective action of tens or even hundreds of thousands of regulatory elements, most of which are annotated as putative enhancers and located in regions far from the promoter regions of genes (Sheffield et al., 2013). Enhancers recruit key TFs and other cofactors to influence the transcription of nearby genes, are usually cell type- and stimulus-specific (Ong and Corces, 2011; Heinz et al., 2015), and play an essential role in brain development and function (Carullo and Day, 2019). While the mapping of open chromatin by DNase/ATAC-seq or the epigenetic landscape (e.g., H3K4me1, H3K27ac) by ChIP-seq have provided a wealth of information about potential enhancers (Ernst et al., 2011), discerning their activity or function in different contexts remains challenging.

To improve our understanding of gene regulation underlying addiction-related behaviors, we profiled the activity of regulatory elements in the brains of rats exhibiting addiction-like behaviors using a recently developed technique called capped small(cs)RNA-seq (Duttke et al., 2019). csRNA-seq captures short initiating (20-60 nt) RNAs with a 5’ cap structure synthesized during the earliest stages of transcription initiation by RNAP II. The method reveals the genome-wide transcription start sites (TSSs) of both stable and unstable transcripts and, thus, all active regulatory elements, including promoters and enhancers, which we will collectively refer to as transcription start regions (TSRs). Since changes in enhancer RNA transcription serve as one of the most reliable markers for nearby gene regulation (Mikhaylichenko et al., 2018), csRNA-seq profiles can provide critical information about the state of regulatory networks in the cell (Duttke et al., 2019; Lim et al., 2021). Furthermore, the single-nucleotide resolution of csRNA-seq data provides a high-resolution mapping of regulatory elements and can reveal spacing relationships between individual transcription start sites (TSS) and TF binding sites (Duttke et al., 2019).

Here, we compared transcription initiation profiles by csRNA-seq using brain tissues isolated from rats that were not exposed to drugs or were subjected to a well-validated extended access model of intravenous self-administration (IVSA) of oxycodone or cocaine (Ahmed and Koob, 1998; Ahmed et al., 2000, 2002; George et al., 2008; Chen et al., 2013; Koob et al., 2014; de Guglielmo et al., 2019; Carrette et al., 2021). Tissues were collected after five weeks of prolonged abstinence to study the long-term effects of voluntary drug intake and were obtained from a tissue repository (Carrette et al., 2021). We selected NAc for its role in mediating the reinforcing effects of substances of abuse and the prefrontal cortex (PFC) for its role in inhibitory control behavior altered in addiction (Everitt, 2014). We integrated active TSR profiles with bulk and single-cell epigenomic data from rat brains to characterize active regulatory elements genome-wide. By comparing drug-exposed versus control samples, we identified potential TFs binding sites differentially transcribed at key enhancer elements in rats with a history of addiction-like behavior. Overall, these findings show the advantage of profiling initiating transcripts to facilitate the identification of upstream regulators of addiction-like phenotypes.

## Materials and Methods

### Brain samples

Brain samples from male heterogeneous stock (HS) rats (2 naive, 2 cocaine, 2 oxycodone) were obtained from the cocaine oxycodone (www.oxycodonebiobank.org) and (www.cocainebiobank.org) tissue repositories at UCSD and are part of an extensive and ongoing study of addiction that uses outbred HS rats (www.ratgenes.org) (Solberg Woods and Palmer, 2019). We selected samples collected during prolonged abstinence after the last session of extended access to oxycodone or cocaine IVSA (Carrette et al., 2021). In this model, male Heterogenous Stock (HS) rats were trained to self-administer drugs in short access conditions (2 h/day for 4 days for oxycodone or 2 h/day for 10 days for cocaine) followed by long access conditions (12 h/day for oxycodone and 6 h /day for cocaine) for 14 days to develop escalation of drug intake. Following the escalation phase, the rats from the oxycodone cohort were characterized for motivation (progressive ratio responding), withdrawal-induced hyperalgesia (mechanical nociception, von Frey test), and development of tolerance to the analgesic effect of opioids (tail immersion test). For the cocaine cohorts, rats were characterized for motivation (progressive ratio responding), compulsive-responding for drug use (contingent footshock), and irritability-like behavior (bottle-brush test). An Addiction Index was computed by integrating all the behavioral measures (Kallupi et al., 2020; Carrette et al., 2021; Sedighim et al., 2021). HS rats classified as having a high Addiction Index were used for this study. Age-matched naive male rats that were not exposed to any drug were used as control. Lastly, brain punches of PFC and NAc tissues were collected after 5 weeks of abstinence. Brain tissue was extracted and snap-frozen (at −30°C). Cryosections of approximately 500 microns were used to dissect PFC and NAc punches on a –20°C frozen stage. Bregma for PFC: 4.20-2.76 mm, and for NAc: 2.28-0.72 mm (3 sections were combined for each).

### csRNA-seq library preparation

We extracted total RNA from PFC and NAc tissues dissected from 6 rat brains using Trizol Reagent (Invitrogen, Cat, num. 15596018) and Zirconium Beads RNase Free (Next Advance, Cat. num. ZrOB05-RNA 0.5 mm) with the Bullet Blender Blue (Next Advance, Model. num. BBX24B) at speed 6 for 1 min. The RNA was purified according to the manufacturer’s instructions (Invitrogen).

csRNA-seq was performed as described previously (Duttke et al., 2019). Briefly, small RNAs of ~15– 60 nt were size selected from 0.3–1.0 microgram of total RNA by denaturing gel electrophoresis. A 10% input sample was taken aside, and the remainder enriched for 5’-capped RNAs.

Monophosphorylated RNAs were selectively degraded by Terminator 5’-phosphate-dependent exonuclease (Lucigen). Subsequent 5’ dephosphorylation by quickCIP (NEB) followed by decapping with RppH (NEB) augments Cap-specific 5’ adapter ligation by T4 RNA ligase 1 (NEB)(Hetzel et al., 2016). Thermostable quickCIP was used instead of rSAP, and hence the bead clean-up step was skipped before heat denaturation before the second round of CIP treatment. The 3’ adapter was ligated using truncated T4 RNA ligase 2 (NEB) before 3’ repair to select against degraded RNA fragments. Following cDNA synthesis, libraries were amplified for 11–14 cycles and sequenced SE75 on the Illumina NextSeq 500 sequencer.

### Hi-C library preparation

One adult SHR/OlaIpcv naive rat was used to generate the Hi-C data. This rat was bred at the University of Tennessee Health Science Center using breeders provided by the Hybrid Rat Diversity Program at the Medical College of Wisconsin. The animal was fully anesthetized by using isoflurane before brains were removed. Brain tissue was extracted and rapidly frozen. Cryosections of approximately 120 microns were obtained in a cryostat set at −11 °C. PFC punches were dissected on a –20°C frozen stage. Tissues were then pulverized in liquid nitrogen. The Arima-Hi-C kit was used to construct the Hi-C libraries (#A410231, Arima Genomics). Sequencing of the libraries was conducted on an Illumina Novaseq S4 instrument by Novogen Inc. The use of rodents was approved by UTHSC IACUC.

### Single nuclei ATAC-seq library preparation

PFC brain tissue from one naive male HS rat was used to generate a single-nuclei ATAC-seq library. Nuclei were isolated from brain tissue as previously described (Corces et al., 2018). Briefly, frozen tissue was homogenized using a 2 ml glass dounce with 1 ml cold 1x Homogenization Buffer (HB). Cell suspension was filtered using a 70 μm Flowmi strainer (BAH136800070, Millipore Sigma) and centrifuged at 350g for 5 min at 4°C. Nuclei were isolated by iodixanol (D1556, Millipore Sigma) density gradient. The nuclei iodixanol solution (25%) was layered on top of 40% and 30% iodixanol solutions. Samples were centrifuged in a swinging bucket centrifuge at 3,000g for 20 min at 4°C. Nuclei were isolated from the 30%-40% interface. Library preparation targeting the capture of ~6000 nuclei was carried out as detailed in the Chromium Next GEM Single Cell ATAC v1.1 manual (10x Genomics). Library sequencing was performed using the Illumina NovaSeq.

### csRNA-seq analysis

Sequencing reads were trimmed for 3’ adapter sequences using HOMER (“homerTools trim −3 AGATCGGAAGAGCACACGTCT -mis 2 -minMatchLength 4 -min 20”) and aligned to the rat mRatBN7.2/rn7 genome assembly using STAR (Dobin et al., 2013) with default parameters. Sequencing statistics are included in **Table S1**. Only reads with a single, unique alignment (MAPQ >=10) were considered in the downstream analysis. Furthermore, reads with spliced or soft clipped alignments were discarded (the latter often removes erroneous alignments from abundant snRNA species). Transcription Start Regions (TSRs), representing 150 bp sized loci with significant transcription initiation activity (i.e. ‘peaks’ in csRNA-seq), were defined using HOMER’s findPeaks tool using the ‘-style tss’ option, which uses short input RNA-seq to eliminate loci with csRNA-seq signal arising from non-initiating, high abundance RNAs that nonetheless are captured and sequenced by the method (full description is available in Duttke et al. (Duttke et al., 2019)). To lessen the impact of outlier samples across the data collected for this study, csRNA-seq samples were first pooled into a single META-experiment per brain tissue region to identify TSRs in each tissue collectively. The resulting TSRs were then quantified in all samples by counting the 5’ ends of reads aligned at each TSR on the correct strand. The raw read count table was then normalized using DESeq2’s rlog variance stabilization method (Love et al., 2014).

The resulting normalized data was used for all downstream analyses. Normalized genome browser visualization tracks were generated using HOMER’s makeMultiWigHub.pl tool (Heinz et al., 2010). TSR genomic DNA extraction, nucleotide frequency analysis relative to the primary TSS, general annotation, and other general analysis tasks were performed using HOMER’s annotatePeaks.pl function. Overlaps between TSRs and other genomic features (including peaks from published studies and annotation to the 5’ promoter using RefSeq defined transcripts), was performed using HOMER’s mergePeaks tool. Functional enrichment analysis of regulated regions was performed using GREAT (McLean et al., 2010) by identifying homologous regions for each TSR in the mouse genome (mm10) using UCSC Genome Browser’s liftOver tool and running GREAT using the mm10 database.

To identify differential TSRs between brain regions or conditions (naive vs. cocaine or oxycodone), we used DESeq2 with FDR < 10% PFC vs. NAc, Naive vs. Oxycodone, Naive vs. Cocaine, or Oxycodone vs. Cocaine.

### Analysis of previously published ChIP-seq and ATAC-seq data

Raw FASTQ files associated with public ChIP-seq and ATAC-seq datasets were downloaded from NCBI’s Short Read Archive and processed in a consistent manner to ensure differences in data processing were minimized for downstream analysis. Reads from ChIP-seq or ATAC-seq datasets were analyzed in a consistent manner. Reads were first trimmed for adapter sequences and then aligned to the rat genome using STAR (Dobin et al., 2013) with default parameters. Only reads with a single, unique alignment (MAPQ >=10) were considered in the downstream analysis. ChIP/ATAC-seq peaks were identified using HOMER’s findPeaks tool using “-style factor” and “-style atac”, respectively. Normalized genome browser tracks were generated using HOMER’s makeMultiWigHub.pl tool. Peak annotations and normalized read density counts were calculated using HOMER’s annotatePeaks.pl tool. Overlapping peaks were determined using HOMER’s mergePeaks.

Datasets used in the study include GR ChIP-seq GSE160806 from the rat hippocampus (Buurstede et al., 2021);. ATAC-seq GSE134935 from rat PFC (Scherma et al., 2020); histone marks and TF ChIP-seqs GSE127793 from rat hippocampal neurons (Brigidi et al., 2019).

### Hi-C Analysis

Hi-C reads were first trimmed for sequences downstream of the restriction/ligation site (“GATCGATC”) and aligned to the rat genome using STAR with default parameters. Normalized interaction contact maps were then generated using HOMER. PCA compartment analysis and topological domain (TAD) calls were generated using HOMER’s runPCAhic.pl and findTADsAndLoops.pl scripts (Heinz et al., 2018). The significant association of A compartment (PC > 1) with ATAC-seq peaks and/or TSRs was calculated using the Mann-Whitney non-parametric Ranksum test.

### DNA motif analysis

Known motif enrichment and *de novo* motif discovery of TSRs were performed using HOMER’s findMotifsGenome.pl tool using 200 bp sequences centered on [-150,+50] relative to TSR primary initiation sites (e.g., strongest TSS in the region) or from −100,100 relative to the center of ATAC-seq peaks (Heinz et al., 2010). When performing *de novo* motif discovery, sequences were compared to a background set of 50,000 random genomic regions matched for overall GC-content. Nucleotide frequency and motif density plots were created using HOMER’s annotatePeaks.pl tool (Heinz et al., 2010). When analyzing ATAC-seq peaks from cell types identified by snATAC-seq, the top 25,000 peaks were selected from each cell type to avoid comparing motif enrichment from sets with large differences in the number of regions that can impact the absolute enrichment levels.

To analyze motif enrichment associated with changes in transcription levels, we analyzed regulated TSRs with MEIRLOP (Delos Santos et al., 2020). Sequences were scored based on their shrunken log2 fold change between treatment conditions (e.g., naive vs. cocaine or oxycodone exposed) and analyzed with MEIRLOP using HOMER’s known transcription factor motif library. The top 3 motifs associated with up- and down-regulation based on their regression coefficients are reported for each comparison (adj. p-values < 0.05).

Furthermore, we provide the BigWig track with the map of transcription factor binding site predictions in the rat genome (rn7), which can be uploaded as a custom track on the UCSC browser as follow:

*track type =bigBed name = “HOMER Known Motifs rn7 (210922)” description= “HOMER Known Motifs rn7 (210922)”*
*bigDataUrl=http://homer.ucsd.edu/homer/data/motifs/homer.KnownMotifs.rn7.210922.bigBedvisibility=3*

### Single nuclei ATAC-seq analysis

Sequencing reads were processed using Cell Ranger ATAC 2.0 with a custom reference for *Rattus norvegicus*, built from the Ensembl Rnor 6.0 release 103 genome and annotation. The filtered results were subsequently analyzed using Signac 1.4.0 (Stuart et al., 2021). Only peaks present in at least 10 cells and cells with at least 200 peaks were considered. Further filtering was performed to retain only cells with between 3,000 and 25,000 fragments, at least 30% of reads in peaks, a blacklist ratio less than 0.05, nucleosome signal less than 4, and TSS enrichment of at least 2.5. Normalization and linear dimensionality reduction were performed using TFIDF, identifying top features with no minimum cutoff and SVD. Nonlinear dimensionality reduction with UMAP and neighbor finding used LSI components 2 through 30, and clustering was performed with the SLM algorithm (Blondel et al., 2008). Cell types were assigned using inferred gene activity. The following cell marker genes were used: *Slc17a* for excitatory neurons, *Gad2* for inhibitory neurons, *Gjai* for astrocytes, *C1qa* for microglia, *Mobp* for oligodendrocytes, *Pdgfra* for oligodendrocytes precursor cells (OPC), *Flt1* for endothelial cells. Pseudo bulk peak positions for each cell type were identified using MACS2 (Zhang et al., 2008). Per-cell TSR enrichment significance was calculated using a one-tailed hypergeometric test and corrected for multiple hypothesis testing using the Bonferroni-Hochberg method.

## Results

### Identification of Transcribed Regulatory Elements in the Rat Brain

To probe if substance abuse can alter gene regulatory programs in the brain, we comprehensively profiled active regulatory elements in two brain regions implicated in addiction: the prefrontal cortex (PFC) and nucleus accumbens (NAc, **Fig. 1A**). Samples from six animals were obtained from a tissue repository (Carrette et al., 2021), including two naive rats, two rats subjected to oxycodone intravenous self-administration (IVSA), and two rats subjected to cocaine IVSA(Arnold et al., 2019; Adhikary et al., 2021; Carrette et al., 2021). We further generated total small RNA-seq libraries that were used as input in csRNA-seq peak calling to mitigate the identification of false TSS from potential RNA degradation-related biases or other high abundance short RNA species. Except for one of the libraries prepared from the NAc of a rat exposed to oxycodone, which failed QC and was discarded from the analysis, csRNA-seq worked as expected by enriching 5’-capped initiating short transcripts (**Table S1, Fig. S1**). As such, the methodological advance of csRNA-seq allowed us to define actively transcribed enhancer RNAs from the banked tissues, which enabled us to explore changes in gene regulatory networks associated with addiction-like behavior.

**Figure 1:**
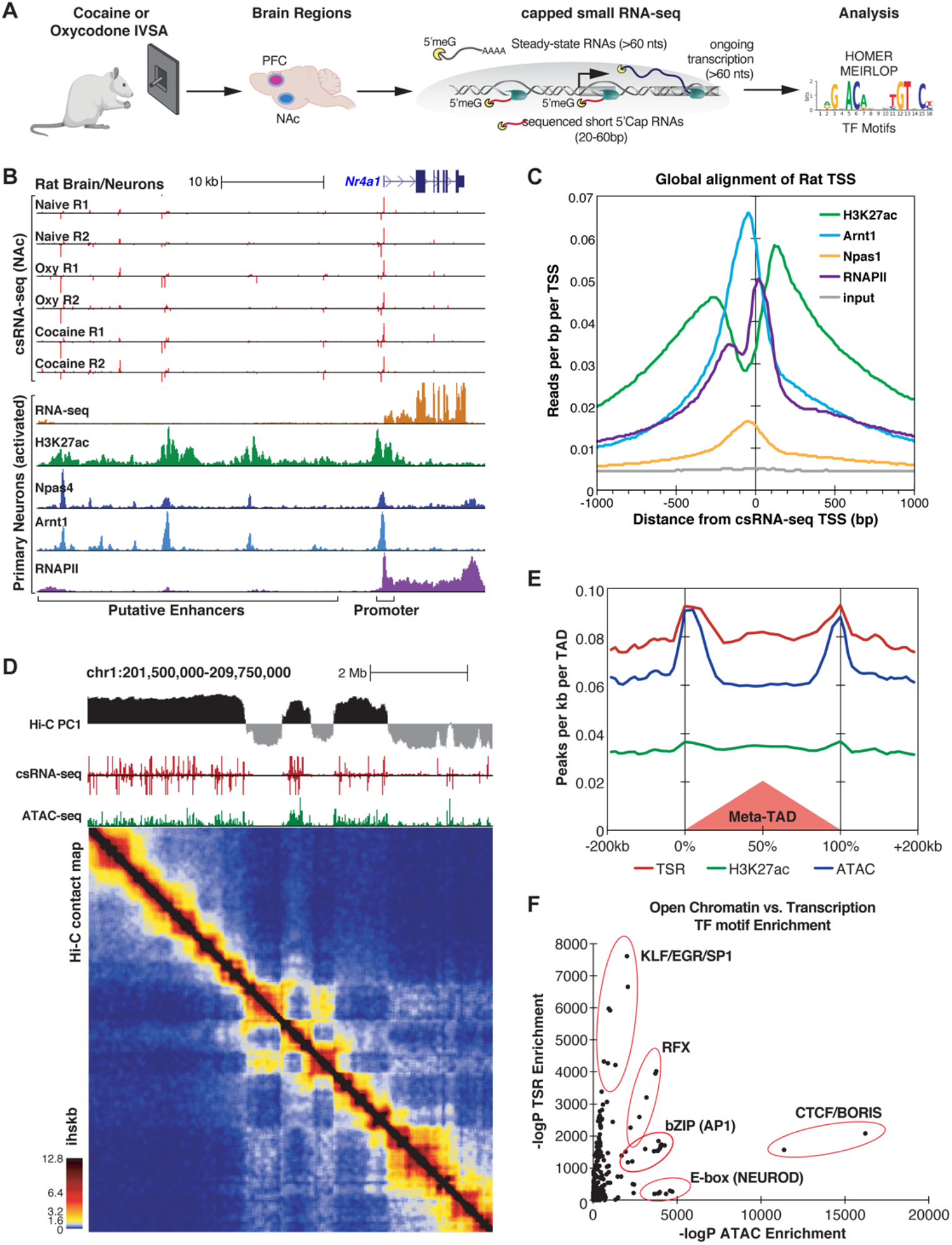
Identification of Transcriptional Start Regions (TSRs) by csRNA-seq in rat brain tissues. **(A)** Diagram of study design. (**B**) An example of csRNA-seq data generated from naive, cocaine-, and oxycodone-exposed rat brains at the *Nr4a1* locus (top) showing overlap with previously published transcriptomic and epi-genomic data from rat hippocampal neurons (bottom). **(C)** Distribution of various histone marks and TFs from primary rat hippocampal neurons with respect to TSRs identified by csRNA-seq in rat brains. Regions are aligned to the primary transcription start site (TSS) in the TSR. **(D)** Genome browser tracks from a representative region of chr1 showing (from top to bottom) A/B chromatin compartments (PC1 from Hi-C), TSRs (csRNA-seq), open chromatin regions (ATAC-seq), and the corresponding contact map of chromatin interactions (Hi-C) from rat PFC tissues. Ihskb = interactions per hundred square kilobases per billion mapped reads. **(E)** Histogram showing the distribution of TSRs, H3K27Ac, and ATAC-seq peaks around TAD regions identified by Hi-C. **(F)** Relationship between ATAC and csRNA motif enrichment for known TF motifs. Motifs recognized by key TFs sharing common DNA binding domains are highlighted.

Across 11 csRNA-seq libraries, we identified 131,647 and 96,563 genomic regions in the PFC and NAc, respectively, with one or more transcription, start sites (TSSs), which we refer to as Transcriptional Start Regions (TSRs). While 15.7% TSRs (20,693 total) in PFC and 19.5% TSRs (18,878 total) in NAc were within or proximate to annotated gene promoter regions, the majority were at promoter-distal sites within introns and intergenic regions of the genome (61% in PFC and 57% in NAc; **Fig. S2A**). These promoter-distal TSRs commonly overlapped with markers of active enhancers from available rat epigenetic data (**Fig. S2B**), as exemplified for the *Nr4a1* locus (**Fig. 1B**). Notably, as seen for the *Nr4a1* locus, distal TSRs were largely bidirectionally transcribed, a common enhancer feature (**Fig. 1B**)(De Santa et al., 2010; Kim et al., 2010; Telese et al., 2015). Analysis of all TSRs genome-wide displayed an architecture typical for vertebrates, with the summit of open chromatin just upstream of the TSSs where the strongest transcription factor ChIP-seq signals can be found (**Fig. 1C**). At the same time, H3K27ac modified nucleosomes were distributed just downstream or upstream of the regulatory region (**Fig. 1C**). Together these data show that csRNA-seq captures active promoters and distal enhancers with high fidelity and accuracy.

The three-dimensional (3D) genome organization can be an essential factor in gene regulation (Andrey et al., 2013; Benabdallah and Bickmore, 2015). To place our identified TSRs in the context of chromatin structure, we generated Hi-C data for the PFC of one rat. 83% of TSRs overlapped with A compartments (PC>0), which define the active region of the genome (**Fig. 1C**) (Lieberman-Aiden et al., 2009). Notably, the association with A compartment was significantly stronger *(p* <1e-16) for transcribed accessible regions (n=91323 ATAC-seq peaks that overlapped a TSR) compared to those that were not transcribed (n=45389 ATAC-seq peaks that did not overlap a TSR, **Fig. S2C**). TSRs also overlapped with the enrichment of H3K27Ac and ATAC-seq peaks at topological domains (TAD) boundaries (**Fig. 1D, E**), which supports the role of promoters and enhancers in defining the boundaries genome-wide (Dixon et al., 2012). Contrasting transcribed and untranscribed open chromatin regions revealed the enrichment of CTCF or helix-loop-helix (bHLH) TFs (e.g., NEUROD1 or OLIG2) in regions with little or no transcription (**Fig. 1F, Fig. S2D**). At the same time, KLF/SP1, RFX, and AP1 motifs were highly enriched in actively transcribed ones (**Fig. 1F, Fig. S2D**), suggesting that these TFs may act as critical activators of brain transcriptional programs. Together, these data emphasize the advantage of capturing enhancer RNAs through methods such as csRNA-seq to define active enhancers over a more basic definition of enhancers simply based on open chromatin or ATAC-seq peaks.

### Brain Region Specificity of TSRs

Enhancers play a critical role in regulating tissue-specific gene expression (Levine, 2010). To identify specific transcriptional signatures for each brain region, we therefore compared TSRs from PFC and NAc, which identified 2,967 PFC-specific and 5,991 NAC-specific TSRs (>2-fold difference, FDR < 10%). Differential TSRs were commonly found near genes typically expressed in the specific brain region. For example, TSRs at the *Neurod6* gene locus were highly transcribed in the PFC but not in the NAc, while the dopamine receptor-1 (*Drd1*) gene locus was highly transcribed in the NAc but not in the PFC (**Fig. 2A**). These results are consistent with the known cellular composition of these brain regions, with the PFC enriched in NEUROD6-expressing glutamatergic excitatory neurons and the NAc enriched in DRD1-expressing medium spiny projection neurons. In addition, this analysis showed that the tissue-specific TSRs are often located adjacent to one another and map within the same TAD (**Fig. 2A)**, suggesting that distal TSRs might preferentially function within a TAD.

**Figure 2:**
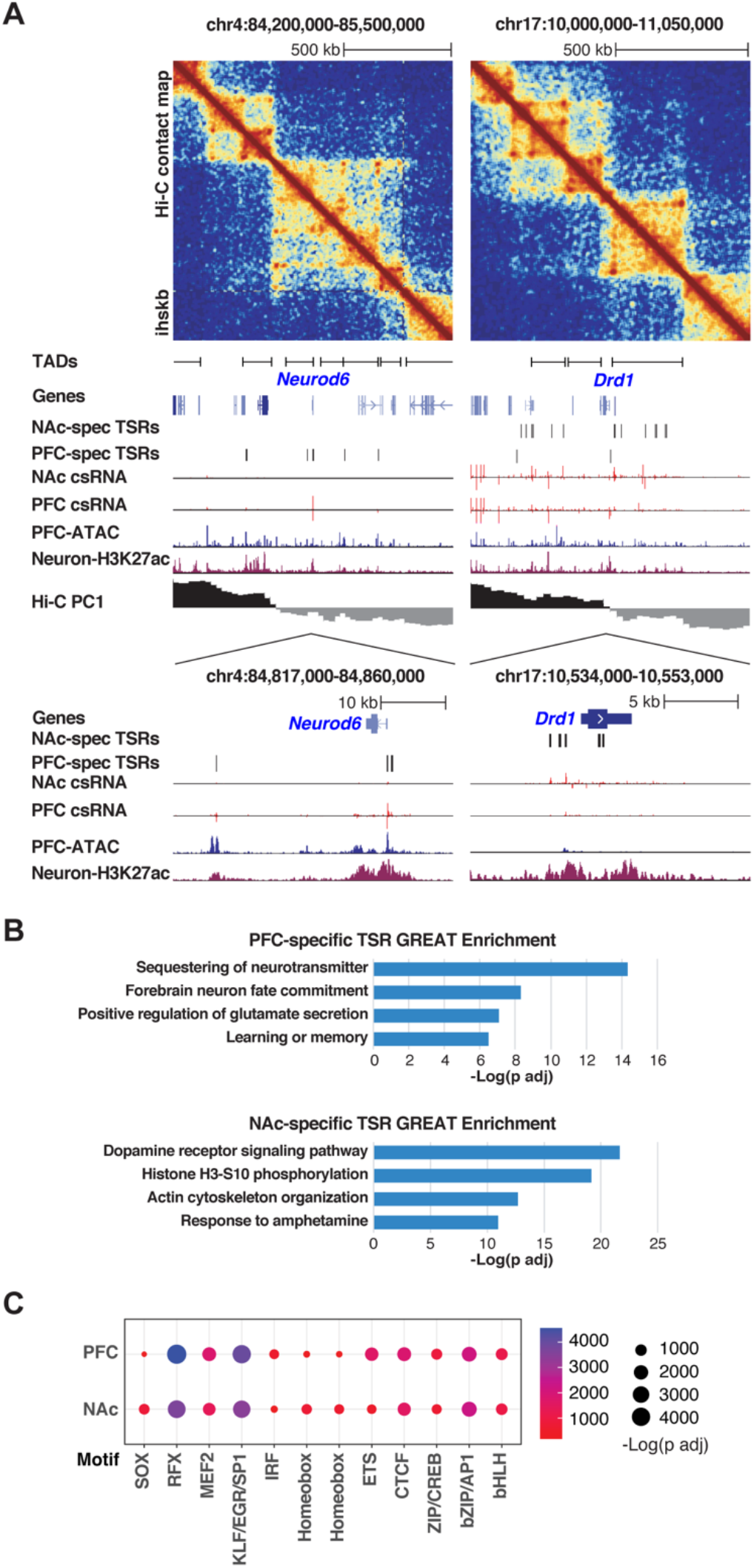
Brain region specificity of Transcriptional Start Site Regions (TSRs). (**A**) *Neurod6* (left) and *Drd1* (right) gene loci are visualized, including (top to bottom) Hi-C contact matrix, TAD positions, genome browser tracks showing tissue-specific TSRs (csRNA-seq), chromatin accessibility (ATAC-seq), active histone mark (H3K27Ac), and A/B compartments (Hi-C PC1). Ihskb = interactions per hundred square kilobases per billion mapped reads. (**B**) Functional annotations associated with the genes near tissuespecific TSRs for PFC (top) and NAc (bottom) as determined by GREAT using mouse genome annotations (see methods). (**C**) Dotplot showing the enrichment scores of known TF motifs in TSRs from PFC and NAc. Size and color of the dots represent the - log adjusted *p*-value as determined by HOMER.

These results were corroborated by pathway analysis of genes found in the vicinity of tissue-specific TSRs. TSRs specifically regulated in PFC were enriched near genes involved in glutamate receptor signaling and learning and memory, supporting the known function of cortical areas in cognitive functions (**Fig. 2B, upper panel**). On the other hand, the TSRs specifically regulated in NAc were enriched near genes in the dopamine receptor signaling pathway and response to psychostimulants, which support the role of NAc in mediating the rewarding effects of substances of abuse (**Fig. 2B, bottom panel**).

The tissue specificity of TSRs was also confirmed by the motif enrichment analysis (**Fig. 2C**). In both regions, TSRs were highly enriched in motifs recognized by general TFs, including the KLF/EGR/SP1 family TFs, basic leucine-zipper (bZIP) TFs (e.g., CREB and AP1 family members) as well as more brain-specific TFs such as MADS-box TFs (e.g., MEF2 family members), and RFX family members (Di Bella et al., 2021; Li et al., 2021; Yao et al., 2021; Zhang et al., 2021; Ziffra et al., 2021). However, these results differed slightly between PFC and NAc. Specifically, PFC-specific TSRs were enriched preferentially for ETS and ISRE motifs, while NAc-specific TSRs were enriched preferentially for RFX, SOX, and Homeobox motifs (**Fig. 2C**).

Taken together, these results show that TSRs profiling from repository tissue is a valid approach to decode tissue-specific regulatory networks, which may be crucial to identify the TFs driving addiction-related transcriptional programs in a brain region-specific manner.

### Comparison of Oxycodone/Cocaine/Naive Rats Reveals Activated and Repressed Regulatory Programs Associated with Addiction-like Behaviors

We next sought to identify regulatory elements associated with a history of addiction-like behavior. Because normalized csRNA-seq read counts across all samples segregated most strongly based on their brain region of origin (**Fig. S3A**), we limited our analysis to comparing conditions within the same brain regions. Using a statistical threshold of > 2-fold difference and FDR < 10%, we identified 317 and 90 differentially regulated TSRs associated with addiction-like behavior in NAc and PFC, respectively (**Fig. 3A, S3B-C**). Notably, oxycodone IVSA resulted in a larger number of differential TSRs than cocaine IVSA in both regions (**Fig. 3A, S3B-C**). Some TSRs were regulated in multiple comparisons, including several near *Hif3a* and *Fkbp5* loci. Moreover, differential TSRs were also enriched near genes that have been previously linked to addiction processes (*Foxo3* (Ferguson et al., 2015), *Tlr4* (Wu and Li, 2020)) or addiction vulnerability (*Nat1* (Comings et al., 2000), *Ppm1k* (Carr et al., 2007; Liang et al., 2010), *Pknox2* (Zuo et al., 2014)).

**Figure 3:**
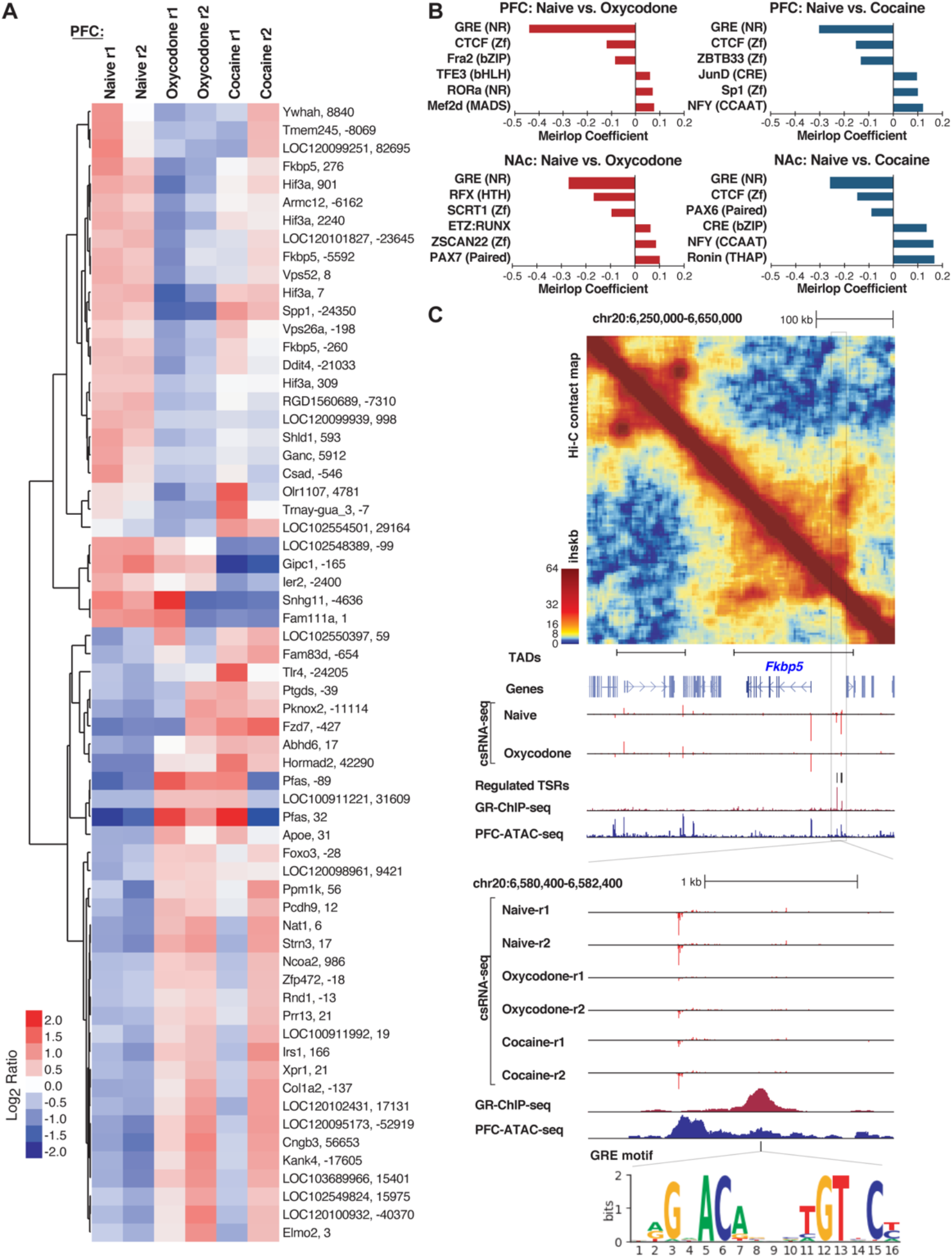
Differentially regulated Transcriptional Start Sites (TSRs) in naïve versus cocaine or oxycodone exposed rat brains. **(A)** Heatmap of transcription initiation levels from differential TSRs in PFC naïve, oxycodone- and cocaine-exposed rats based on mean-centered log2 ratios; each row shows the closest gene and the TSR position relative to that gene’s annotated TSS. **(B)** Barplot of significant logistic regression MEIRLOP coefficients for topranked motifs associated with regulated TSRs between naïve and oxycodone or cocaine conditions in PFC and NAc. **(C)** Example of regulation at the *Fkbp5* gene locus, including (top to bottom) Hi-C contact matrix with TAD positions, genome browser tracks showing regulated TSRs (csRNA-seq), GR binding (ChIP-seq), chromatin accessibility (ATAC-seq), and GRE motif location. Ihskb = interactions per hundred square kilobases per billion mapped reads.

To gain insights into the TFs that may mediate changes in gene expression networks in response to a history of substance abuse, we identified TF motifs enriched in TSRs regulated by oxycodone or cocaine exposure in each brain region. To do so, we used MEIRLOP (Brigidi et al., 2019; Delos Santos et al., 2020), a DNA motif analysis approach that associates motifs with the magnitude of regulation at TSRs across conditions based on a logistic regression. This analysis identified a strong and consistent association between the glucocorticoid response element (GRE) and TSRs down-regulated in brain tissue from rats with addiction-like phenotypes versus controls (**Fig. 3B**). Our identification of GRE-binding TFs as potential key regulators of addiction-related reprogramming of gene regulatory networks is consistent with the well-established role of glucocorticoid signaling in addiction (Srinivasan et al., 2013; Koob et al., 2014). Furthermore, our analysis identified bZIP motifs for AP1 family members (e.g., CREB, JUN, FOS) as enriched in TSRs up-regulated in both brain regions from rats exposed to cocaine compared to naive rats (**Fig. 3B**), which is consistent with previous findings showing activation of members of the AP1 family in addiction-related processes, such as ΔFOSB or CREB (Teague and Nestler, 2021).

To validate the motif enrichment predictions, we next overlapped regulated TSRs with GR binding sites previously identified in the rat hippocampal neurons (Buurstede et al., 2021). We found that 12 of the 32 TSRs down-regulated in oxycodone-exposed PFC were within 1 kb of a GR ChIP-seq peak (*p* < 0.0002). To further support GR’s potential role in regulating these TSRs, several downregulated TSRs were found in the intergenic region upstream of *Fkbp5* (**Fig. 3C**), a well-known GR target gene. Analysis of Hi-C data in this region identified a TAD that encompasses the *Fkbp5* locus and includes the cluster of regulated TSRs associated with addiction-like behavior (**Fig. 3C**), which provides evidence for GR binding and GRE motifs in the nearby regulatory DNA.

Together, the unbiased discovery of TSRs, combined with motif analysis, uncovered TF-driven gene regulatory programs associated with addiction-like phenotypes in rats.

### Cell Type Specificity of TSRs Associated with Addiction-like Behaviors

Enhancers often function in a highly cell type-specific manner (Levine, 2010). Understanding the specific cell types of the brain in which enhancers are active may be critical to unlocking important regulatory mechanisms underlying addiction-like behavior. To this aim, we used a cell type-specific reference of chromatin accessibility sites that we generated by snATAC-seq using the PFC of a naive rat (**Fig. 4A**). First, we annotated different classes of brain cell types based on the chromatin accessibility of known cell markers, including excitatory and inhibitory neurons, astrocytes, oligodendrocytes, oligodendrocytes precursor cells, microglia, and endothelial cells, showing that this dataset successfully captured known cell types of the rat PFC. In support of this result, motif enrichment analysis with HOMER (**Fig. 4B**) showed that motifs for lineage-specific TFs are enriched in their expected cell types (e.g., AP1/MEF2C/TBR1 in neurons, PU.1 in microglia, SOX10 in oligodendrocytes). By cross-referencing TSRs with the snATAC-seq, we assigned expressed genes and their active regulatory elements identified by csRNA-seq to specific cell types (**Fig. 4C)**. For example, regulatory elements at gene loci of known cell type-specific markers (e.g., *Olig2, Ctss, Slc32a1, Neurod6)* showed accessible chromatin exclusively in the expected cell types that directly overlapped TSRs identified in the bulk csRNA-seq experiments (**Fig. 4B**).

**Figure 4.**
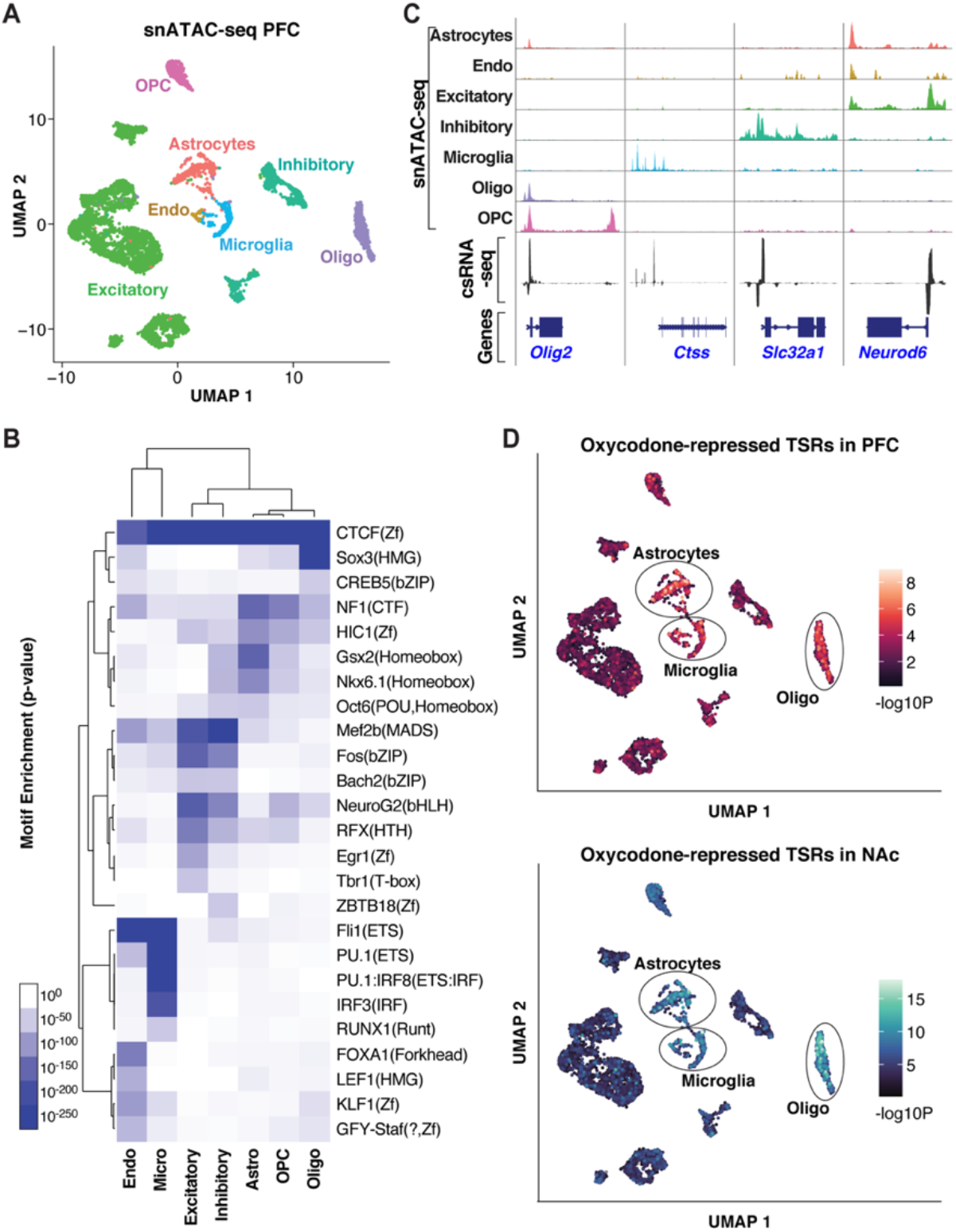
Cell-type assignment of active regulatory elements (TSRs). **(A)** UMAP clustering of cells based on snATAC-seq of the PFC. Clusters are colored based on cell types inferred from the accessibility patterns near known marker genes. **(B)** Genome browser tracks of pseudo bulk ATAC-seq read densities showing genes with cell-type-specific snATAC-seq profiles and csRNA-seq from bulk tissue. **(C)** TF motif enrichment across accessible regions from specific cell types in the snATAC-seq data. **(D)** UMAP visualization of oxycodone-associated repressed TSRs enriched in individual cells identified by snATAC-zseq in PFC and NAc, showing consistent enrichment in astrocyte, microglia, and oligodendrocyte populations

To address the cell-type specificity of the gene regulatory networks associated with addiction-like behavior, we sought to map the addiction-regulated TSRs to the different cell types identified by snATAC-seq. To this aim, we analyzed the oxycodone-repressed TSRs in the PFC and NAc, which were strongly enriched in GRE motifs (**Fig. 3B**). This analysis revealed that the downregulated TSRs overlapped accessible regions enriched in non-neuronal cells, such as astrocytes, microglia, and oligodendrocytes (**Fig. 4D**), suggesting the involvement of glial cells in the regulatory programs underlying addiction-like behaviors. Given that the repressed TSR were enriched in GRE motifs, this result also suggests a role of GR in regulating transcriptional responses to opioids, specifically in glial cells.

These results highlight the advantage of integrating csRNA-seq with snATAC-seq data to probe the cellular specificity of gene regulatory mechanisms and highlight the role of glial cells in modulating addiction-related behavior.

## Discussion

Here we report the active transcriptional landscape of the PFC and NAc from rats with a history of addiction-like behaviors. By integrating transcriptional initiation (csRNA-seq) with genome structure (HiC) and single-cell epigenomic data (snATAC-seq), the analysis of the regulatory landscape not only provided a comprehensive catalog of eRNAs but also identified TFs that are likely to play important regulatory roles. Using this approach, we discovered that GR-bound enhancers are strongly down-regulated during prolonged abstinence from oxycodone or cocaine IVSA, specifically in glial cells.

There is strong evidence supporting the role of cell type-or stimulus-specific enhancers in the gene regulation (Heinz et al., 2015), but determining whether an enhancer is active in specific cellular or biological states remains a significant challenge in the field. Recent studies using nascent transcriptional profiling suggest that the transcriptional states of enhancers are better predictors of active chromatin states than open chromatin or histone modifications (Danko et al., 2021). However, many nascent transcriptional methods have technical limitations, including the requirement of intact nuclei and large numbers of cells. csRNA-seq overcomes these limitations by quantifying the level of transcription initiation at regulatory elements, such as enhancers, from total RNA, which can be easily obtained from frozen tissues (e.g., samples from a tissue repository). Using csRNA-seq on < 1 μg of total RNA isolated from repository brain tissues, we identified > 100k TSRs across PFC and NAc from naive rats or rats with addiction-like behavior following oxycodone or cocaine IVSA (Carrette et al., 2021). Most TSRs represent eRNAs as they initiate transcripts in regions associated with known features of enhancer elements, including open chromatin, histones harboring the H3K27ac mark, and bidirectional transcription. Although the function of eRNAs is still controversial(Li et al., 2016), converging lines of evidence show that their abundance is highly correlated with the expression of proximal genes and precedes stimulus-dependent transcription of the mRNA of these genes (Kaikkonen et al., 2013; Arnold et al., 2019). Thus, identifying active enhancers is likely important to decipher the gene regulatory basis of addiction. Furthermore, combining csRNA-seq with TF motif discovery provides different and complementary information than traditional transcriptomic or epigenetic data (e.g., ATAC-seq). As such, it can be used as an unbiased functional assay for TF activity.

The major finding of this study is the identification of TF-regulatory networks associated with a history of addiction-like behavior. The analysis of drug-altered TSRs revealed that GR-regulated enhancers were consistently repressed in PFC and NAc from rats with a history of oxycodone and cocaine addiction-like behavior compared to controls. This result is consistent with converging evidence that the brain stress system involving glucocorticoid signaling plays a critical role in the development of addiction-like behavior (George and Koob, 2010; Koob et al., 2014). The cell-type deconvolution analysis also showed that repressed TSRs in the PFC and NAc were enriched in glial cells, which is consistent with findings suggesting that alterations of neuroimmune mechanisms such as neuroinflammation or synaptic remodeling by glial cells can contribute to the liability of addiction (Lacagnina et al., 2017). Furthermore, a recent single-cell transcriptomic study found a robust transcriptional response to acute morphine treatment in oligodendrocytes and astrocytes of the mouse NAc (Avey et al., 2018). Several morphine-induced genes identified in this study were GR targets, supporting a role of GR in regulating transcriptional responses to opioids. In line with this notion, GR has been shown to modulate opioid reward processing by regulating genes essential for astrocytic metabolism (Slezak et al., 2013; Skupio et al., 2020). However, our results show an opposite direction of transcriptional regulation that the different treatment protocols may explain (acute versus chronic exposure), or it may reflect negative feedback mechanisms of glucocorticoid signaling during stress responses associated with addiction-related phenotypes (prolonged abstinence vs. acute withdrawal)(Srinivasan et al., 2013). It is also important to note that our results do not entirely preclude the involvement of different TFs that recognize similar motifs, including mineralocorticoid, androgen, or progesterone receptors. Further experiments targeting GR or its targets in specific celltypes of rodent models of addiction will be necessary to validate the role of GR in different addiction-like behaviors.

Our study has several limitations. First, we used a limited number of samples (n=2/condition), which may lead to a low statistical power to detect differentially expressed TSRs and could explain why, despite identifying over 100,000 TSRs across two brain regions, we only detected a relatively small number of differentially regulated TSRs in both PFC and NAc. A study with a larger cohort of rats would be ideal for confirmation. Secondly, the control animals used in this study are rats that were never exposed to drugs; thus, our study design does not consider environmental factors associated with the behavioral protocol (e.g., surgery, foot-shock, pharmacokinetics factors). Including rats with a low addiction index subjected to the same behavioral protocol but do not develop addiction-like phenotypes would serve as important control to provide more substantial evidence that the differences we observe reflect molecular changes associated with addiction-related processes rather than other phenomena. Lastly, our study only includes male rats, which precludes the analysis of sex differences in regulatory networks associated with the known sexual dimorphism of addiction-like behaviors (Fattore and Melis, 2016).

In summary, we used an unbiased and highly sensitive method to identify active enhancers by measuring levels of initiating transcripts from brain tissues of rats with addiction-like phenotypes. We identified TF-centered regulatory mechanisms implicated in addiction, including those regulated by GR in glial cells. Overall, our study demonstrates that transcriptional initiation profiling has the potential to dissect the gene regulatory mechanisms driving addiction-related phenotypes in an unbiased and quantitative manner.

## Supplementary Figures and Legends

**Supplementary Figure 1:**
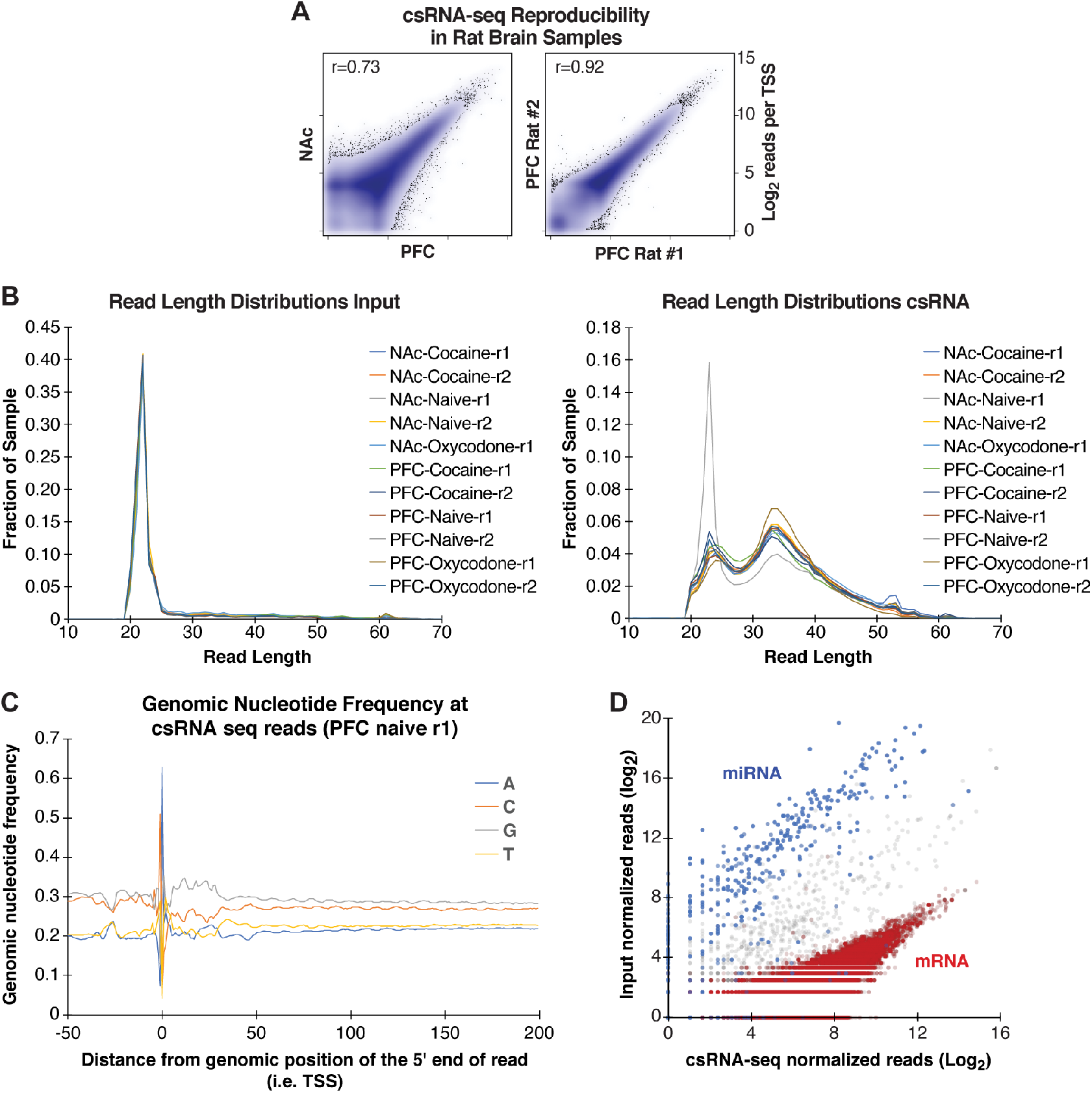
csRNA-seq data in rat brain tissues. **(A)** Variation in csRNA-seq levels at each Transcriptional Start Site Region (TSR) between tissues (NAc vs. PFC) or between replicates (PFC r1 vs. r2) in samples from naive rat brains. (**B)** Read length distribution for input libraries (left) and csRNA-seq libraries (right). Input libraries show a strong spike at 21 nt corresponding to mature miRNA. (**C)** Nucleotide frequencies at csRNA-seq reads shown for the PFC naive-r1 library. (**D**) Read counts at the annotated promoters (5’ end of transcripts -/+ 200bp) with blue dots indicating miRNA transcripts, red dots mRNA transcripts, and grey dots other transcripts (snRNAs, snoRNAs, etc.). csRNA-seq and input data corresponding to the PFC naive-r1 sample is shown.

**Supplementary Figure 2:**
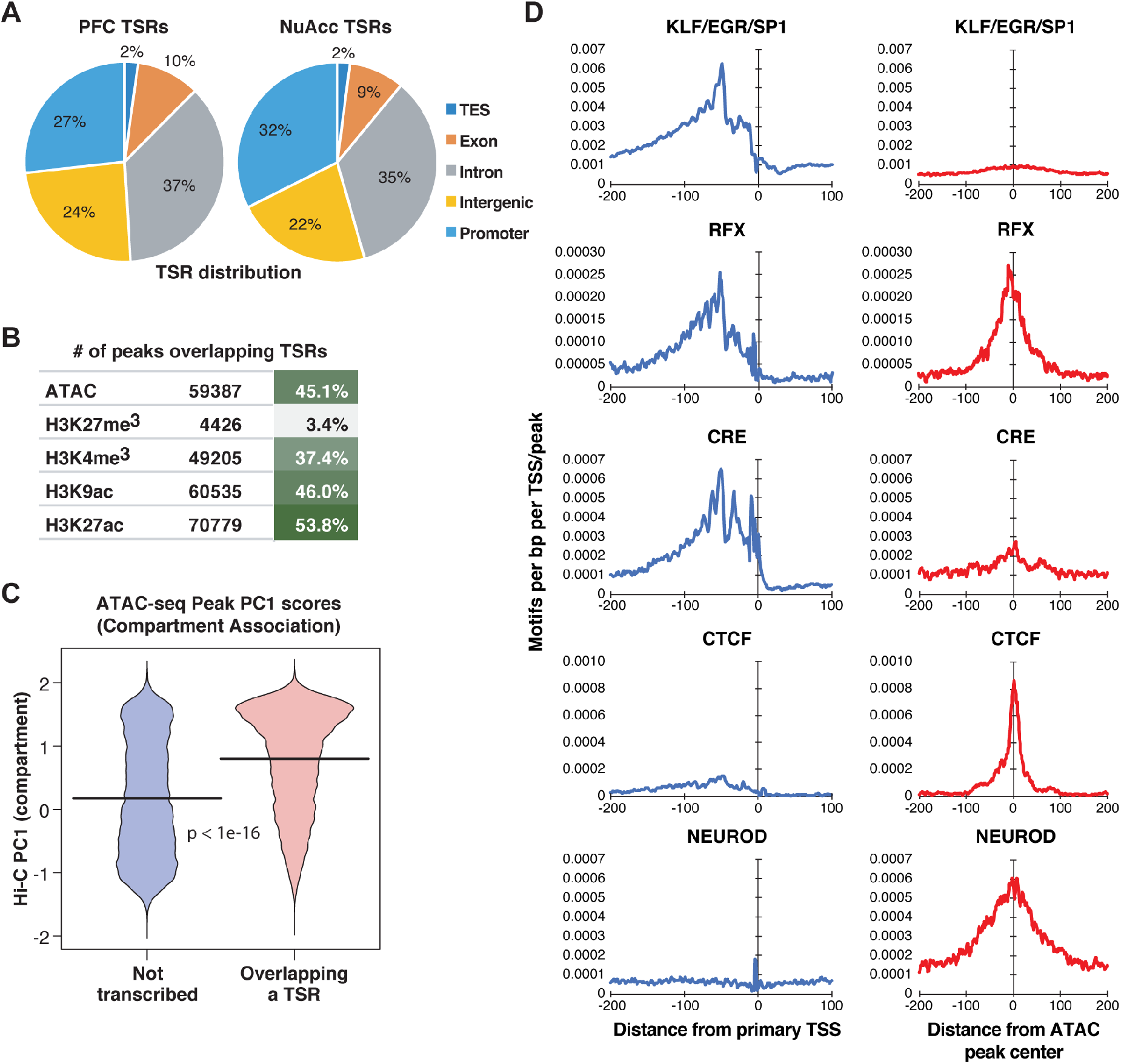
Identification and characterization of Transcriptional Start Sites (TSRs) from csRNA-seq data. **(A)** Pie charts showing the distribution of TSRs in different genomic regions. (**B**) Fraction (%) of TSRs that overlap peaks identified from ATAC-seq or ChIP-seq for several histone marks. The number of peaks analyzed is reported. (**C)** Violin plot showing the distribution of Hi-C PC1 values for ATAC-seq peaks that are not transcribed or overlapping a TSR. **(D)** Location of several TFs motifs with respect to the primary TSS from csRNA-seq TSRs and the center of ATAC peaks genome-wide.

**Supplementary Figure 3:**
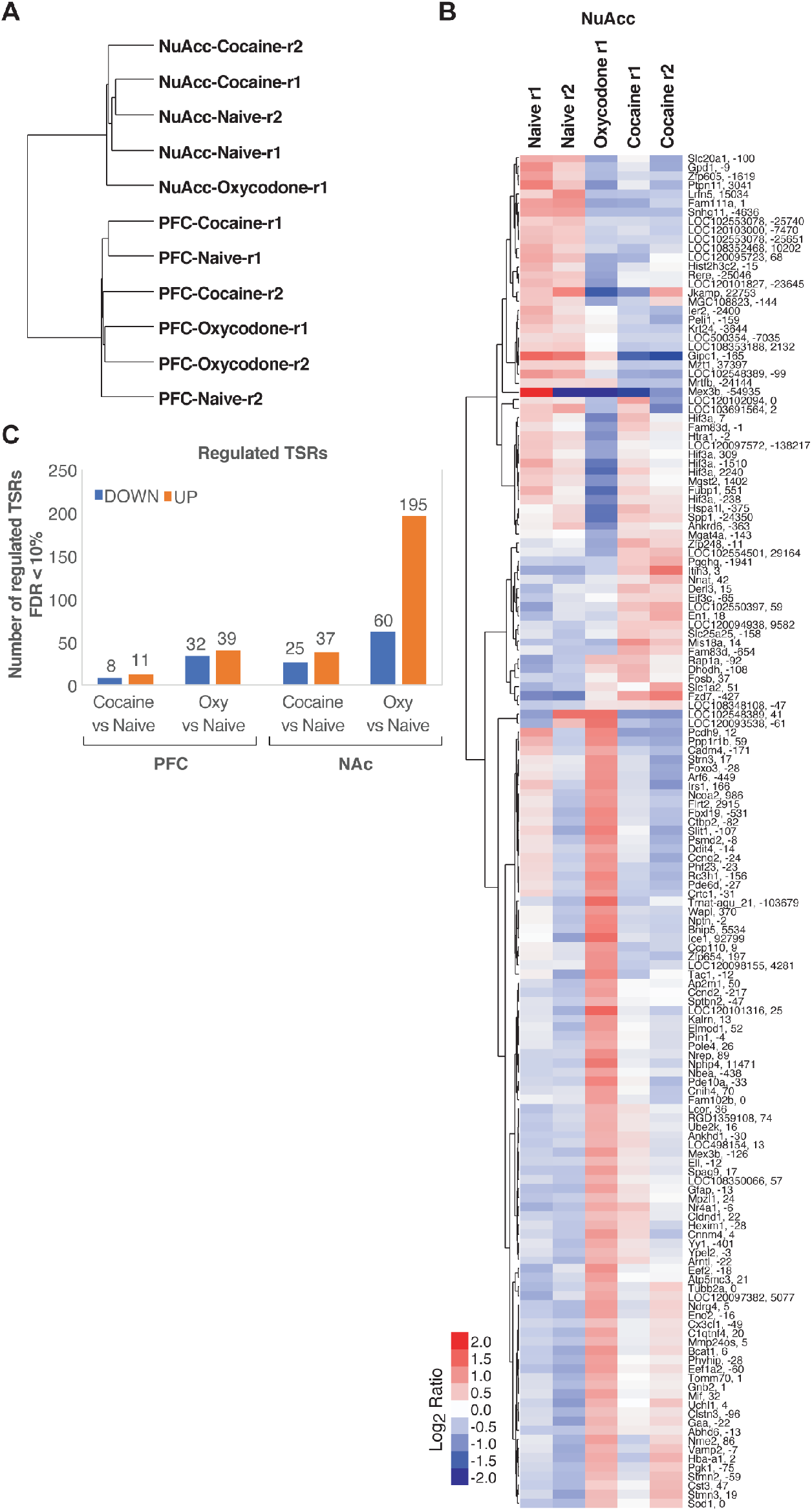
Differentially regulated Transcriptional Start Sites (TSRs) in naïve versus cocaine- or oxycodone-exposed rat brains. **(A)** Hierarchical clustering of csRNA-seq samples shows segregation based on brain regions. (**B**) Heatmap of differential TSRs in NAc naïve, oxycodone- and cocaine-exposed rats based on log2 ratios relative to the mean; each row showing the closest gene, TSR position relative to TSS, and chromosomal coordinates. (**C**) Number of differentially regulated TSRs (>2 fold, FDR < 10%) across naïve and drug-exposed conditions.

## Acknowledgments

We want to thank Lisa Maturin for their technical assistance. This work was supported by NIH (K99GM135515 to S.H.D.; DA050239 to F.T.; DA051972 to F.T. and C.B.; GM134366 to C.B; DA053672 to H.C.; P50DA037844 to A.A.P.; DA043799, DA037844 to O.G. This publication includes data generated at the UC San Diego IGM Genomics Center utilizing an Illumina NovaSeq 6000 that was purchased with funding from a National Institutes of Health SIG grant (#S10 OD026929).

